# Hybridization and a mixture of small and large-effect loci facilitate adaptive radiation

**DOI:** 10.1101/2022.02.18.481029

**Authors:** Rishi De-Kayne, Oliver M. Selz, David A. Marques, David Frei, Ole Seehausen, Philine G. D. Feulner

## Abstract

Adaptive radiations represent some of the most remarkable explosions of diversification across the tree of life. However, the constraints to rapid diversification and how they are sometimes overcome, particularly the relative roles of genetic architecture and hybridization, remain unclear. Here, we address these questions in the Alpine whitefish radiation, using a whole-genome dataset that includes multiple individuals of each of the 22 species belonging to six ecologically distinct ecomorph classes across several lake-systems. We reveal that repeated ecological and morphological diversification along a common environmental axis is associated with both genome-wide allele frequency shifts and a specific, larger effect, locus, associated with the gene *edar*. Additionally, we highlight the role of introgression between species from different lake-systems in facilitating the evolution and persistence of species with unique phenotypic combinations and ecology. These results highlight the role of both genome architecture and secondary contact with hybridization in fuelling adaptive radiation.

## Introduction

Understanding the genetic basis of speciation and adaptive radiation without geographic isolation, and determining how and when such diversification is possible, is a key aim of evolutionary biology. Speciation with gene flow often occurs in the form of ecological speciation. During this process, reproductive isolation results from divergent ecological selection, or ecologically-mediated, sexual selection ^1,2^. Despite the supposed prevalence of ecological speciation in adaptive radiation, the factors that constrain or facilitate speciation and the mechanisms by which speciation proceeds during the adaptive radiation of lineages are still not well understood ^3,4^. Studying the identity and genomic distribution of loci involved in ecological speciation, particularly in cases where parallel ecomorphological contrasts have arisen multiple times, is one way to address these questions.

Using such approaches, studies have already highlighted the prevalence of either strongly differentiated genomic ‘islands’ of differentiation ^5–10^ or genome-wide polygenic architectures of phenotypic differentiation and ecological speciation ^11–15^. Both of these architectures on their own constrain speciation with gene flow in different ways. In the former scenario, reproductive isolation may be unlikely to evolve because the chance that loci under divergent selection will be linked to a trait that causes reproductive isolation is slim, and a genome-wide correlated response to divergent selection is lacking ^16,17^. Polymorphism may therefore be a more likely outcome than speciation. In the polygenic scenario, the strength of per-locus divergent selection is likely to be small and insufficient to overcome the homogenising effects of gene flow ^18^. Combinations of a larger number of genome-wide small-effect loci and some large-effect loci, may therefore be most conducive for overcoming constraints to speciation in the face of gene flow that result from either one of these architectures alone ^4,7,19^.

In addition to specific genetic architectures, empirical ^20–24^, experimental ^25,26^, and theoretical ^27,28^ investigations have implicated introgression between non-sister species as a process that may also facilitate diversification and adaptive radiation. Since introgression generates novel combinations of haplotypes, combining those from the distinct parental species, it may be possible that hybrid populations are able to span fitness valleys and, in turn, occupy ecological niches that would otherwise be inaccessible via stepwise adaptation ^27^. However, few studies have been able to link ecological novelty with empirical signatures of introgression in the wild (but see ^22,29^).

The Alpine whitefish radiation contains over 30 species of the genus *Coregonus*, which have evolved in small species flocks across multiple lake-systems in the last 10-15 thousand years ^30–36^. Although whitefish have speciated in many postglacial lakes across the Northern hemisphere, species flocks in pre-Alpine lakes are particularly diverse, and up to six whitefish species, with different ecological strategies, exist in sympatry, and exhibit considerable phenotypic variation, including body size, spawning depth and season, gill-raker count and length, and diet (Fig. 1a; ^30–33,36^). Across multiple lake-systems, species from different species flocks have evolved similar ecological strategies and phenotypes and as such have been categorised into ecomorphs based on these similarities ^37^. Interestingly, a number of traits are correlated across the Alpine whitefish radiation, particularly amongst widespread ecomorphs that have evolved in most species flocks. For example, deeper spawning species tend to have higher gill-raker counts and smaller bodies than shallower spawning species (Fig. 1b; Supplementary Fig. 1; ^38^). However, in addition to these widespread ecomorph trends, there are a number of less-widespread ecomorphs that have evolved just in one or few lakes and exhibit different trait combinations, decoupled from this trend (for example deep spawning, small bodied, species with few gill rakers). The fact that sympatric whitefish species flocks are thought to have evolved independently within each lake-system ^35^ provides the opportunity to identify overarching genomic features that may have facilitated rapid diversification, including the rapid and repeated evolution of similar ecomorphs, and the origin and persistence of species with new trait combinations.

**Figure 1.**
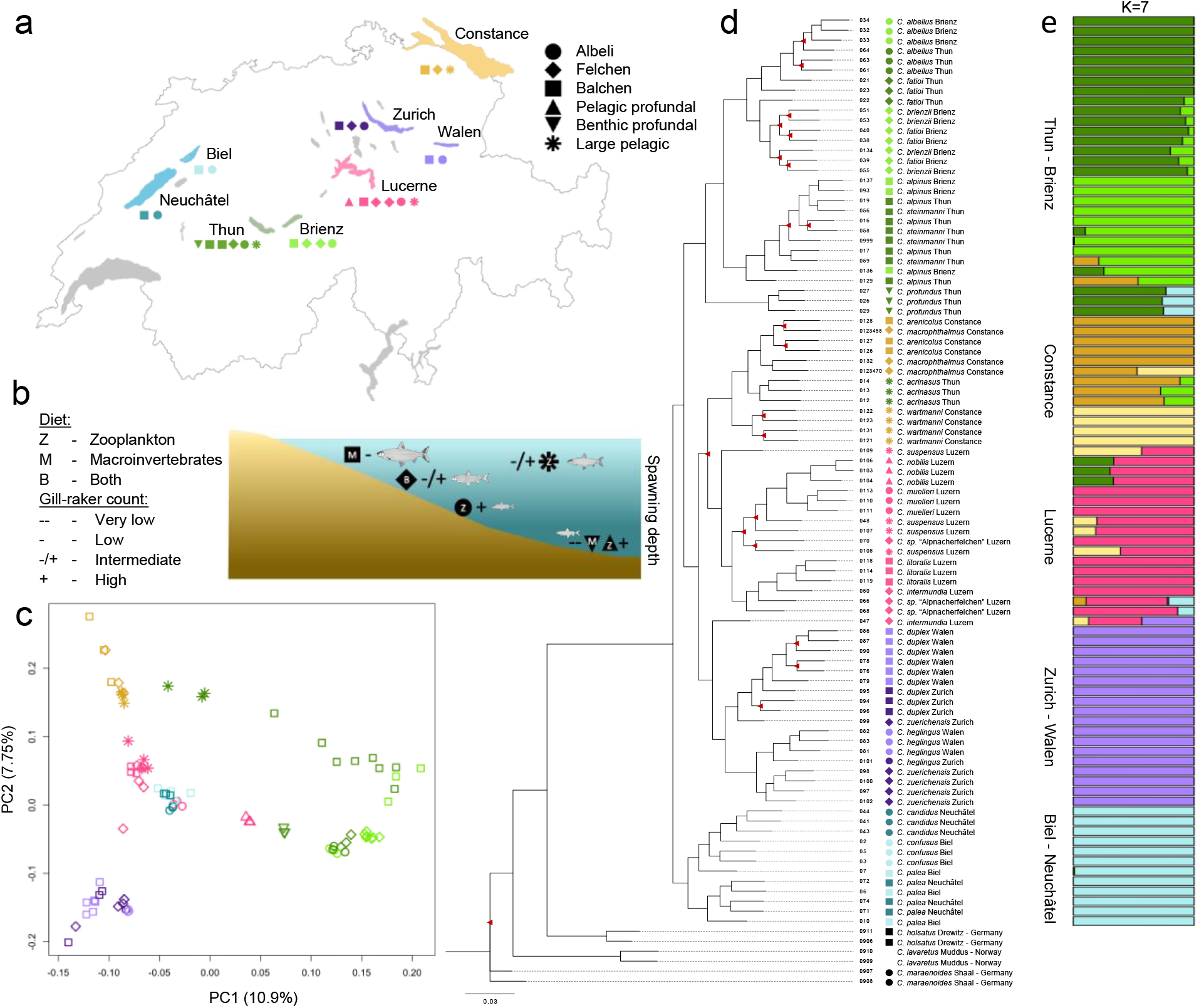
Lake-systems and species are genomically distinct across the Alpine whitefish radiation. a) A map of the Alpine whitefish species, assigned to ecomorphs, sampled from each of the pre-Alpine lake-systems including Constance (yellow), Zurich/Walen (purple), Lucerne (pink), Thun/Brienz (green) and Biel/Neuchâtel (blue). b) A qualitative diagram showing the ecological characteristics of each whitefish ecomorph (represented by different symbols) including relative spawning depth (indicated by position in figure), diet (indicated by letter), and relative gill-raker count (indicated by −/+ symbols); fish illustrations by Verena Kälin. c) A genomic PCA of all 91 Alpine whitefish based on a linkage-disequilibrium filtered SNP-set of 1,133,255 (a subset of our full 14,313,952 SNP dataset) which separates out the Thun/Brienz system from all other lakes on PC1 and each of the other lake-systems from one another on PC2. d) A maximum likelihood RAxML phylogeny produced using a thinned subset of 1,692,559 SNPs from all 99 sequenced whitefish individuals (nodes have bootstrap support ≥ 95/100 unless highlighted with red triangles; outgroup samples with known ecomorph assignment are denoted with black symbols; for ease of viewing the most distantly related outgroup C. albula is pruned from this tree). e) An admixture analysis highlights the lake-system based population structure within the Alpine whitefish radiation, and shows that sympatric whitefish species are each other’s closest relatives (to best observe within and between-lake-system level population structure, K=7 is shown; see Supplementary Fig. 2 for the range of CV error associated with other values of K and Supplementary Fig. 3 for admixture proportions of individuals from K=2 to K=10)

Here, we build on past work on adaptive radiations ^39,40^ to investigate the genetic basis of diversification, and the ways in which, in the absence of geographical isolation, constraints to speciation may have been overcome, within the Alpine whitefish radiation. We compiled whole-genome sequences for 99 whitefish individuals, spanning 22 species belonging to six distinct ecomorphs, from five pre-Alpine lake-systems (putative species flocks), and four outgroup species (Fig. 1a; Fig. 1b; Supplementary File S1). We show that phenotypic diversification along water depth gradients, which is independently repeated across five lake-systems ^41^, is underpinned by a mixed genetic architecture that comprises both genome-wide differentiation and one locus with a larger effect on phenotype. Additionally, our results suggest that secondary contact between species from different species flocks, followed by interspecific hybridization has helped overcome constraints to the evolution of additional niche specialists.

## Results

### Phylogeny and population structure

To understand how the Alpine whitefish radiation evolved and how the relationships between sympatric species within flocks, and between ecologically similar species (belonging to the same ecomorph) in different flocks, are structured, we produced a genomic PCA (Fig. 1c) and constructed a phylogenetic tree (Fig. 1d). Our PCA and phylogeny confirm and expand upon results of earlier work ^35^ in demonstrating that the Alpine whitefish radiation is monophyletic with respect to non-Alpine whitefish and European Cisco (*Coregonus albula;* not plotted), and, in general, each of the pre-Alpine lake-systems sampled constitutes a reciprocally monophyletic species flock. Both the branching patterns in the phylogeny and the results of our clustering analysis (Fig. 1d; Fig. 1e; K=7; Supplementary Fig. 2; Supplementary Fig. 3) are concordant with the independent evolution of sympatric species flocks within lakes or lake-systems, and hence the parallel evolution of species with similar ecological strategies, i.e. ecomorphs. The one substantial deviation from this pattern of reciprocal monophyly amongst lake-system species flocks is the placement of *C. acrinasus*, which phylogenetically belongs to the Lake Constance clade despite being endemic to Lake Thun (discussed below; also noted in ^35^; in addition to a number of individuals with putative hybrid signatures).

### Parallel allele frequency shifts underpin repeated ecological differentiation

Of the six whitefish ecomorph classes, the most widely distributed are the large, deep bodied, and macro-invertivorous ‘Balchen’, the smaller, shallower bodied, and zooplanktivorous ‘Albeli’, and the ‘Felchen’, which have intermediate characteristics between these two ecomorphs (across these three widespread ecomorphs, whitefish species exhibit correlated trait variation; Fig. 1b; Supplementary Fig. 1). Our phylogeny indicates that within each lake, two genetically distinct lineages typically emerged first, separating a ‘Balchen’ species from an ‘Albeli’ species or, if ‘Felchen’ species are present, from the common ancestor of ‘Albeli’ species and ‘Felchen’ species (with the exception of Lake Constance where no ‘Albeli’ species is present). These divergence events therefore happened separately in each lake-system, and species belonging to these widespread ecomorphs evolved independently in different lake-systems. To identify whether this parallel phenotypic differentiation was underpinned by parallel allele-frequency shifts we first investigated four sympatric pairs of ‘Balchen’ and ‘Albeli’ species from lakes Brienz, Lucerne, Walen, and Neuchâtel. We subsetted our full data set to include three ‘Balchen’ and three ‘Albeli’ individuals from each of these four lakes and first analysed F4 statistics to confirm that indeed each sympatric species-pair represents a single independently evolved species-pair (as in ^42^). Topologies placing sympatric ‘Balchen’ and ‘Albeli’ species as sister taxa in a four-taxon tree had consistently lower F4 statistics, indicative of a more accurate topology, than topologies where the species of the same ecomorph from different lakes were sister taxa (Supplementary Fig. 4). Then we calculated the cluster separation score (CSS) between the ecomorph groups (i.e. individuals of the four ‘Balchen’ species were grouped together and individuals of the four ‘Albeli’ species were grouped together; ^43,44^; Fig. 2a), allowing the detection of signals of parallel allele frequency differences between ecomorphs. The resulting 1659 50 kb CSS outlier windows, which represented parallel allele frequency shifts between the ‘Balchen’ and ‘Albeli’ species from different lakes (identified by running a permutation test which shuffled the assignment of individuals to each ecomorph group whilst maintaining population structure and then identifying windows with an FDR corrected p-value of < 0.01), were distributed genome-wide (Fig. 2a). These 1659 parallel-differentiated windows overlapped with 1800 genes in total, which were significantly enriched for a number of gene ontology terms including those related to neurons, cell signalling, and fatty acid metabolism (Supplementary File S2 contains a full list of significantly enriched gene ontology terms; Supplementary Fig. 5 shows that the length distribution of these genes was not substantially different to that of all annotated genes).

**Figure 2.**
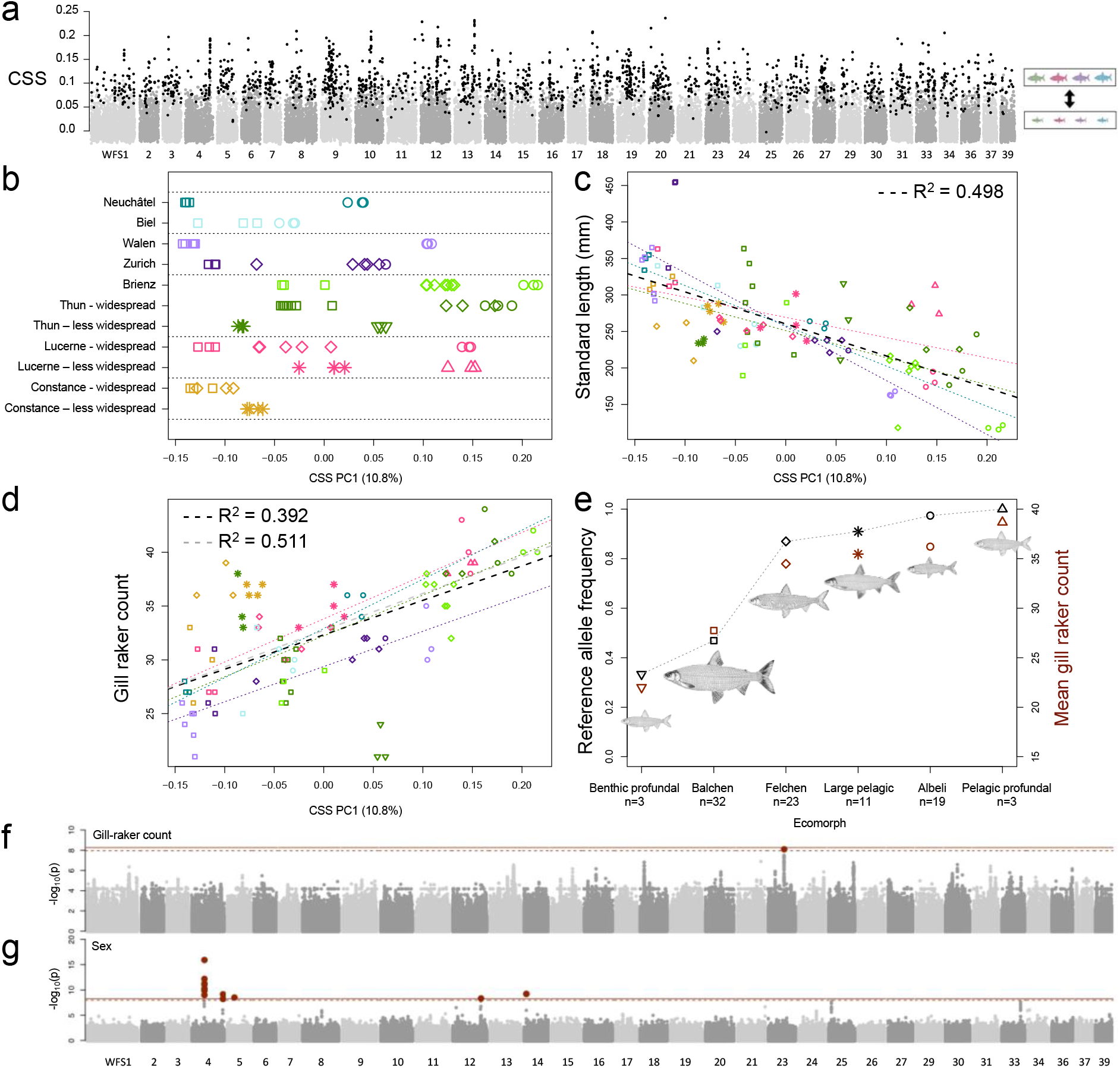
A combination of genome-wide allelic variation and major effect loci underpins parallel ecomorph differentiation in the Alpine whitefish radiation. a) CSS scan (calculated between 12 ‘Balchen’ individuals and 12 ‘Albeli ‘ individuals, with each ecomorph group comprising four distantly related species) highlights genome-wide parallel allele frequency shifts between ‘Balchen’ and ‘Albeli ‘ ecomorphs across the four lakes. Outlier CSS windows are shown in black. b) PC1 calculated using linkage filtered SNPs from across the 1659 CSS outlier windows for all whitefish individuals separates whitefish within lakes and lake-systems (rows separated by dashed lines). Widespread and less-widespread ecomorphs within the same lake are separated along the same axis. c) Whitefish standard length plotted against CSS PC1 for all lakes together (black line; R^2^=0.498, p=8.06×10^-15^) and for each lake separately. Significant lake-system-specific regressions are coloured by lake-system and range in R^2^ from to 0.322 in Lake Lucerne (p=0.01405) to 0.6925 in the lake Walen/Zurich system (p=1.84×l0^-5^). d) Gill-raker count plotted against CSS PC1 for all lakes together including (black line; R^2^=0.3921, p=4.1×10^-11^) or excluding (grey line; R^2^=0.5107, p=7.63×10^-15^) the outlier species C. profundus, and for each lake-system separately. Significant lake-system-specific regressions are coloured by lake and range in R^2^ from 0.3871 in the Lake Thun-Brienz system (including the outlier C. profundus; p=1.11×10^-4^; when excluding C. profundus R^2^=0.6051, p=4.22×10^-7^) to 0.8113 in lake Lucerne (p=3.47×10^-7^). See Supplementary Table S1 for all details regarding lake-specific statistics. e) Allele frequencies for the SNP significantly associated with gill-raker count variation where all 91 Alpine whitefish are grouped by ecomorph (black symbols) compared to ecomorph-averaged gill-raker counts (red symbols); fish illustrations by Verena Kälin. f) GWAS results for gill-raker count and g) sex for all 9,120,498 polymorphic SNPs within the Alpine whitefish radiation across the 90 individuals with corresponding phenotypes.

Genetic variation across CSS outlier regions not only differentiated ‘Balchen’ and ‘Albeli’ species from each other but also allowed the separation of species belonging to the four other whitefish ecomorphs within each lake-system (Fig. 2b; Supplementary Fig. 6). We further show that genomic variation across these parallel differentiated regions (captured by CSS PC1; Fig. 2b) correlated with body size (standard length; Fig. 2c; total R^2^=0.498, p= 8.06×10^-15^; see Supplementary Table S1 for lake-system-specific statistics) and gill-raker count (Fig. 2d; total R^2^=0.3921, p=4.1×10^-11^; see Supplementary Table S1 for lake-system-specific statistics), suggesting that in addition to explaining variation between ‘Balchen’ and ‘Albeli’ species, these genomic regions might contribute to broader phenotypic differences between other ecomorphs, including intermediate ‘Felchen’ species and to some degree the less-widespread ecomorphs, ‘large-pelagic’, ‘benthic-profundal’, and ‘pelagic-profundal. These results are concordant with a scenario of polygenic differentiation between sympatric species, with many loci affected by divergent selection and potentially associated with ecological and phenotypic differences and each contributing a small amount to a broader overall pattern of divergence.

### Parallelism in gene functional pathways between independent ecomorph contrasts

In addition to patterns of genetic parallelism between species of the widespread ‘Balchen’ and ‘Albeli’ ecomorphs, we also investigated each of the four independently evolved ‘Balchen’ and ‘Albeli’ species pairs separately, to identify whether, despite the presence of parallel allele frequency shifts, the most strongly differentiated genomic regions between ecomorphs are species-pair-specific or shared among replicate pairs from different lakes. Species-pair-specific patterns of strong differentiation may be indicative of subtle differences in selection regimes between lakes and hint at the degree to which genetic redundancy, where different genotypes can result in similar phenotypes, underpins parallel ecomorph differentiation. As such, we assessed whether genomic differentiation between each independently evolved ‘Balchen’ and ‘Albeli’ species-pair involved the same set of alleles, genes, or gene pathways, hinting at the commonality of ecomorph evolution across lake-systems. To understand the genome-wide landscape of differentiation across the four independent ‘Balchen’ and ‘Albeli’ species pairs we first carried out separate pairwise F_ST_ scans in 50 kb windows (each with >10 SNPs) for each sympatric species-pair (resulting in ~34,000 windows for each species-pair; Supplementary Fig. 7). This window-based approach averaging F_ST_ estimates based on only 12 alleles across multiple loci may result in some observed frequency differences arising from sampling, limiting us to the detection of strong selection and near fixation regimes ^45,46^ but allows us to explore the degree of genomic redundancy across scales. The most differentiated regions of the genome between sympatric ‘Balchen’ and ‘Albeli’ species (outlier windows within the top F_ST_ percentile for a given species-pair) have a genome-wide distribution (with mean genome-wide background FST across the four species pairs ranging from 0.06 in Neuchâtel to 0.12 in Brienz; Supplementary Fig. 7), and are species-pair-specific, with no outlier windows shared across all four lakes (6 outlier windows were shared between three contrasts, and 63 shared between two; in keeping with findings from North American whitefish ecomorph contrasts where observed genetic differentiation is not parallel across all lakes ^47^). These species-pair-specific patterns were also reflected at the gene level (i.e. regardless of window boundaries), where, out of 1130 genes that overlapped with F_ST_ outlier windows in at least one of the four sympatric ‘Balchen’ and ‘Albeli’ species contrasts (out of the 42,695 genes that sit on scaffolds that were annotated in the reference genome), none overlapped with an outlier window in all four lakes (Supplementary Table S2).

The lack of overlap in genes associated with outlier windows across the four species pairs may also suggest that genetic redundancy is at play. To test whether genetic redundancy explains species-pair-specific differentiation patterns we investigated whether the same set of four species pairs exhibit parallelism at the functional level rather than at the gene level by comparing gene orthology terms and pathways associated with each gene that overlapped with FST outlier windows between sympatric ‘Balchen’ and ‘Albeli’ species. For the 1130 genes overlapping F_ST_ outlier windows we identified 660 KEGG ortholog terms, of which two were associated with outlier windows in the species pairs of all four lakes (Supplementary Table S2). For both of these orthology terms from the KEGG orthology database that were associated with outlier windows in all four lakes (K07526 and K12959) we found that in Lake Neuchâtel one associated gene was on chromosome WFS12, and in the remaining three lakes a second associated gene was located on chromosome WFS10 (K07526 is also associated with an additional gene in Lake Lucerne). For K07526, both genes, despite being located on different chromosomes, had BLAST hits to different isoforms of the protein *SRGAP3* (SLIT-ROBO Rho GTPase-activating protein 3). Similarly, for K12959 both genes hit to caveolin and caveolin-like proteins in other salmonids. WFS12 and WFS10 are homeologous chromosomes ^48^, supporting the idea that genomic redundancy, in this case across homeologous chromosomes, is involved in ecomorph differentiation. This finding furthermore supports the idea that the ancient salmonid-specific whole-genome duplication facilitated diversification by increasing the number of possible adaptive combinations of alleles ^49^. Additionally, around one third (111/ 315) of the KEGG pathways that the 660 KEGG ortholog terms belonged to were associated with outlier windows in all four independent species pairs (Supplementary Table S2). This shared differentiation at the metabolic pathway level, across independent speciation events with similar phenotypic outcomes, without parallelism at the gene level, highlights the role of genetic redundancy. As such, parallel ecomorphological divergence across the radiation may be underpinned by a polygenic adaptive architecture featuring redundancy, as reflected by the many parallel frequency shifts detected (using CSS), the lack of widely shared regions of strong differentiation (as indicated by F_ST_), and the evidence for genetic redundancy at the gene pathway level ^50^.

### Large-effect loci underpin a key ecological trait

We also identify the genetic basis of variation in gill-raker count in whitefish, a key ecological trait that often differs between species occupying different niches because of its role in determining feeding efficiency on different prey items, i.e. trait utility ^51,52^. Fish with fewer gill-rakers feed most efficiently on benthic macroinvertebrates ^53^ whilst fish with many gill-rakers feed most efficiently on zooplankton ^51^. We tested associations between gill-raker counts and SNPs (those polymorphic within the Alpine whitefish radiation). Using data from all 90 Alpine whitefish individuals with recorded gill-raker counts we identified a single significantly associated SNP on WFS23 (−log10(p)=8.1; LD-considerate significance threshold −log10(p)=7.96; Fig. 2f), that explains 31% of the variation observed in gill-raker counts and displays highly correlated allele frequencies with mean gill-raker counts across all ecomorphs and species (Fig. 2e; Supplementary Fig. 8). This candidate SNP fell within an annotated whitefish gene on WFS23, which, when aligned with other salmonid assemblies using BLAST, hit with high confidence against the *edar* gene (ectodysplasin-A receptor). This gene is known to be involved in gill-raker development in zebrafish, where *edar* knockouts exhibit a loss of gill-rakers ^54^, and is in the same protein family as the gene *eda*, which is known to underpin a number of ecologically important features in other fish species, most notably plating in stickleback ^55^. Using a similar approach, we also identified a number of significant sex-associated peaks, with the most significantly associated SNP (−log10(p)=15.93), explaining 54% of the variation in sex across the radiation, located on WFS04 (Fig. 2g; see methods for more information).

### Hybridization facilitates ecological diversification

Although species of the geographically widespread ‘Balchen’, ‘Felchen’ and ‘Albeli’ ecomorphs repeatedly diverge from one another along the common ecological axis of water depth with correlated phenotypic differentiation in several traits (including standard length and gill-raker count; Fig. 1b; Supplementary Fig. 1), likely the result of similar selection pressures along water depth gradients in different lakes, some lakes additionally harbour species of less-widespread ecomorphs, with distinctive ecological strategies. These include ‘large-pelagic’, ‘benthic-profundal’, and ‘pelagic-profundal’ species. These species have combinations of traits that contrast with the direction of correlation among traits seen in the widespread ecomorphs. For example, whereas species that spawn deeper typically have higher gill-raker counts, reflective of the transition from feeding on benthic macroinvertebrates to zooplankton, the ‘benthic-profundal’ *C. profundus* spawns very deep but has very few gill-rakers. Interestingly, our admixture analysis highlighted that a number of species that belong to these less-widespread ecomorphs, including two of the three ‘large-pelagic’ species, and both profundal species, show evidence of genetic admixture between species flocks from different lakes (Fig. 1e). To investigate these signals further, and determine whether secondary contact and introgression were associated with the evolution and maintenance of less-widespread ecomorphs with distinct trait combinations, explaining their heterogeneous distribution across the Alpine whitefish radiation, we calculated excess allele sharing between species across our dataset. Excess allele-sharing was computed using the f-branch statistic *f_b_*(*C*), which was calculated from f4 admixture ratios, f(A,B;C,O), for all combinations of species (or clades in cases where sister species belong to the same ecomorph but are not reciprocally monophyletic) within and between lakes that fit the relationships ((A, B), C), according to our phylogeny (Fig. 1d).

When considering the three ‘large-pelagic’ species (*C. wartmanni* in Lakes Constance, *C. acrinasus* in Lake Thun, and *C. suspensus* in Lake Lucerne), the most striking significant introgression (indicated by a high, and significant, *f_b_*(*C*) value) reflects excess allele sharing between Lake Constance and *C. suspensus* from Lake Lucerne, particularly with the Constance ‘large-pelagic’ species *C. wartmanni* (Fig. 3; black box). This result is concordant with our admixture analysis which indicated that *C. suspensus* indeed looks admixed between species of Lake Lucerne and Lake Constance. The Lucerne ‘large-pelagic’ *C. suspensus* also appears to have significant, but less substantial, excess allele sharing with a number of other Lucerne species. Our results also suggest, as supported by our phylogeny and admixture analysis, that the ‘large-pelagic’ species in Lake Thun, *C. acrinasus*, is genetically admixed, with alleles from Lake Constance and Lake Thun (indicated by significant excess allele sharing with all Brienz/Thun branches in our tree; Fig. 3). This also confirms, and clarifies, the results of other studies which suggested that the evolution of *C. acrinasus* involved the historical anthropogenic translocation of fish from Lake Constance into Lake Thun ^38^. Despite this extensive gene flow in the recent past, *C. acrinasus* now appears to persist as a stabilised hybrid species, demonstrated by its monophyly in our phylogeny (Fig. 1d) and distinct placement in our PCA (Fig. 1c). Together, these patterns suggest that the ‘large-pelagic’ ecomorph may have originally evolved in Lake Constance, and that fish of this species from Lake Constance subsequently colonised, or were translocated to other lake-systems where hybridization with native species then occurred and hybrid species became established (as suggested by historical records for lakes Thun ^38^ and Lucerne ^56^).

**Figure 3.**
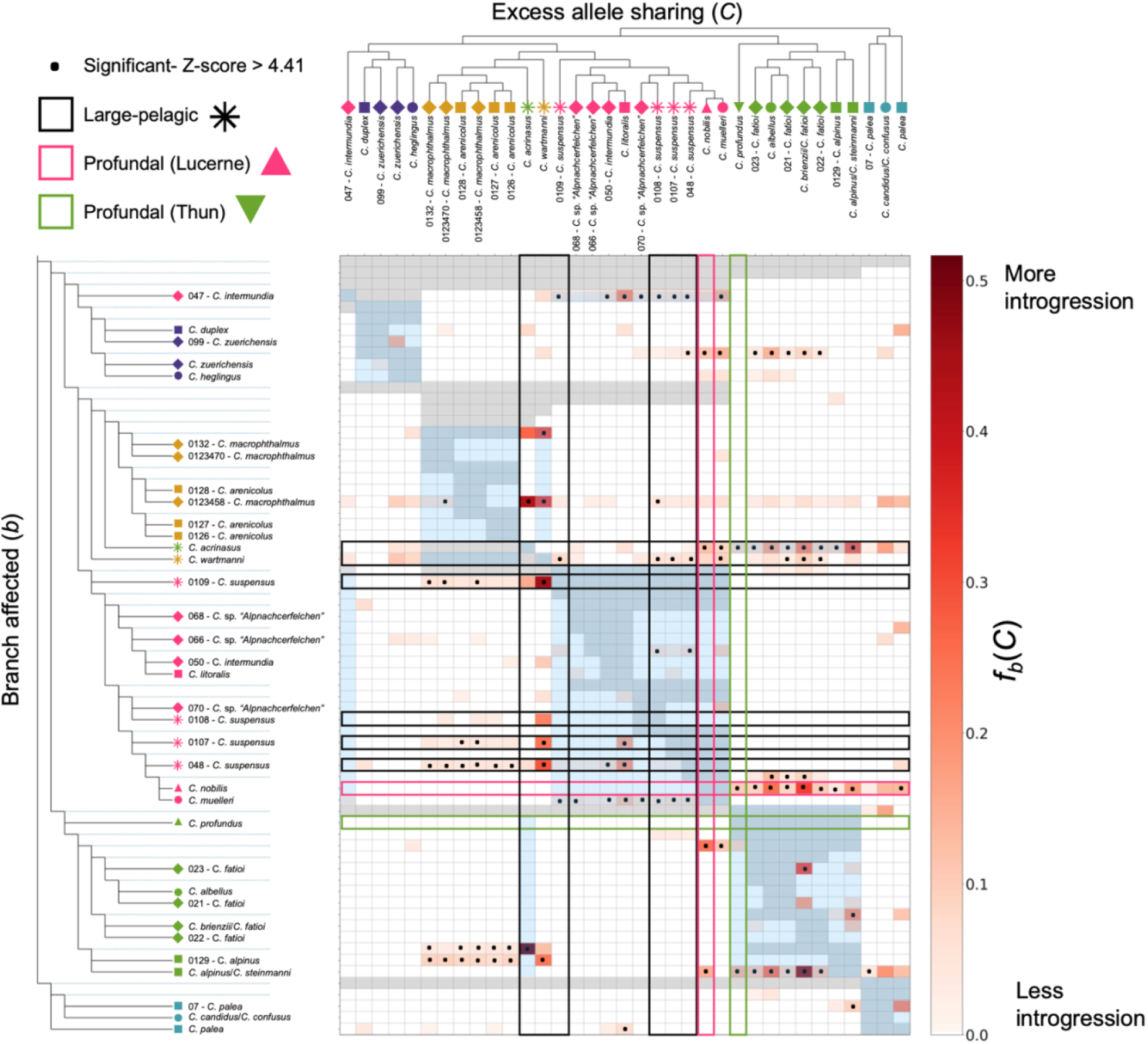
Excess allele sharing is widespread between whitefish species both within and between lake-systems. F-branch f_b_(C)) statistics across our dataset highlight excess allele sharing between tips in the tree (which represent species or individuals when species were not monophyletic; horizontally arranged at the top of the figure) and each other tip and node in the phylogenetic tree (vertically arranged on the left of the figure), compared to its sister branch. The associated lake and ecomorph of each tree tip is indicated by the symbol and colour (as in Fig. 1a). The redness of each cell in the matrix indicates the degree of excess allele sharing between each tree tip (C) and each tip or node (b) with significant instances of excess allele sharing, where the Z-score was >4.41 (equivalent to the Bonferroni multiple testing corrected p-value of 0.01), are highlighted with a dot. For clarity, when a species within a lake or lake-system is supported as monophyletic we have collapsed all of its individuals into a single tree tip. Blue shading is used to indicate comparisons among species within a lake-system. Grey shading indicates tests which cannot be carried out due to the topology of the tree. F-branch statistics associated with species of the three focal ecomorphs are highlighted with boxes in the matrix including the large-pelagic ecomorph of which we have three species from three lake-systems (black), pelagic-profundal ecomorph as a single species from Lucerne (pink) and the benthic-profundal ecomorph as a single species from Thun (green).

Interestingly, a modest amount of excess allele sharing was observed between the ‘benthic-profundal’ species from Lake Thun *C. profundus* and the ‘large-pelagic’ species *C. acrinasus* (from Lake Thun; likely the result of within-lake gene-flow), and the ‘pelagic-profundal’ species *C. nobilis* from Lake Lucerne (Fig. 3; green box), despite the implied admixture from the Biel/Neuchâtel system (as was shown in Fig. 1e). However, more substantial signals of excess allele sharing were observed between other non-profundal ecomorphs of Lakes Thun/Brienz and *C. nobilis* (Fig. 3; pink box). The strongest signals of excess allele sharing with *C. nobilis* came from the ‘Albeli’ species *C. albellus* in lakes Thun and Brienz, and the ‘Felchen’ species from Lake Brienz. The ‘pelagic-profundal’ *C. nobilis* may therefore constitute a stabilised hybrid between other whitefish species from Lake Lucerne and some from Lakes Thun or Brienz. These system-wide f-branch statistics highlight that significant signals of excess allele sharing are less commonly associated with species of widespread ecomorphs which exhibit correlated traits, but are prevalent when considering species of less-widespread ecomorphs, which have trait combinations that are discordant with these correlations.

## Discussion

Adaptive radiations provide a valuable opportunity to identify constraints of diversification and to disentangle the ways in which species may overcome some of these constraints. In this study we addressed these outstanding questions using radiation-wide whole-genome sampling. We found that the genetic basis of rapid parallel evolution of widespread Alpine whitefish ecomorphs comprises both a locus of large effect, implicating the gene *edar* in underpinning gill-raker variation, and many allele frequency shifts distributed across the length of the genome. We were also able to detect parallelism in gene-pathways differentiating species of the ecologically contrasting and widespread ‘Balchen’ and ‘Albeli’ ecomorphs across different lake-systems. Our data also suggest that the evolution and maintenance of less-widespread ecological strategies and unique trait combinations, is often associated with introgression upon secondary contact between species of different species flocks.

Previous empirical and theoretical work had suggested that the genetic basis of differentiation of ecologically contrasting species can comprise few large effect loci ^5–10^ or many genome-wide small-effect loci ^11–15^. However, mounting empirical evidence suggests that polygenic architectures with a combination of these two, a so-called ‘mixed’ genetic architecture comprising many small-effect loci and a few large-effect loci, may also be present ^14,19,57^. Such ‘mixed’ architectures may provide the ideal substrate for rapid speciation in the absence of geographical isolation, since they may better facilitate the build up of linkage disequilibrium in the face of gene flow than either very few key loci or highly polygenic architectures. This is because large-effect loci can act as ‘visible’ targets for selection, and additional genome-wide modifier loci increase the chances of the accumulation of reproductive isolation via linked selection ^18^. Our data suggests that such an architecture, a combination of large and small effect loci, indeed underpins variation among species in the Alpine whitefish radiation.

Our results also highlight the potential role of genetic redundancy in facilitating the repeated evolution of ecologically similar species within adaptive radiations. Genetic redundancy can act at many scales and describes the scenario in which various alleles both within and between genes, and even gene pathways, result in similar phenotypes ^50,58^. Such genetic redundancy may help explain rapid and repeated instances of evolution, since subtly different environment-specific selection regimes acting on different regions of the genome can still drive parallel phenotypic change. It may be possible that the prevalence of duplicated genes (ohnologs) after whole genome duplication, and the possible relaxation of selection acting on these ohnologs ^59^, may facilitate both the *de novo* evolution of novel alleles (and thus phenotypes) and increase the likelihood that different populations can evolve and reach the same fitness optimum in a genetically non-parallel but redundant way. Whole genome duplication is thought to have facilitated adaptation in a diverse array of clades (including plants ^60^, fungi ^61^, and animals ^62^), and our observations that different ohnologs underpin differentiation between ecologically similar, independent, whitefish species pairs support the idea that the ancient salmonid-specific whole-genome duplication facilitated diversification by increasing the number of possible adaptive combinations of alleles (Macqueen & Johnston, 2014).

Whilst our data shows that highly replicated ecomorphological differentiation along similar ecological (water depth) gradients in different lake-systems is underpinned by a mixed genetic architecture, hybridization upon secondary contact between species from different lake-systems seems to facilitate the additional growth of species flocks through addition of species with trait combinations that are decoupled from those associated with speciation on depth gradients. Whilst a mixed genetic architecture promotes the rapid and repeated diversification of ecologically similar whitefish, there are likely constraints to the phenotypic divergence that can be achieved simply by the shuffling of existing alleles. As a result, the occupation of vacant niches may require new combinations of alleles that result in new, discordant, combinations of traits. Hybridization between distantly related species, e.g. non-sister whitefish species from different species flocks, results in the coming together of adaptive alleles or haplotypes which have each been tested by selection on their own, but have not previously existed in these combinations. This gene flow upon secondary contact between separate species flocks within a single large radiation may therefore provide a mechanism by which constraints to diversification may be overcome, allowing evolution into new niche space without having to persist through low-fitness intermediate states ^27,63^. The specific genetic architecture of introgressed regions might also play a crucial role in determining the potential to overcome constraints, since large introgressed haplotypes can rapidly reach substantial frequencies following hybridization ^64^. Our results suggest that a combination of the genetic architecture of traits under divergent selection and the opportunity for secondary contact and hybridization between non-sister species are both important for rapid adaptive radiation.

## Methods

### Sampling the radiation

To understand the phylogenetic relationships between Alpine whitefish we carried out whole-genome resequencing on 96 previously collected whitefish (with associated phenotypic measurements including standard length and gill-raker counts; collected in accordance with permits issued by the cantons of Zurich (ZH128/15), Bern (BE68/15), and Lucerne (LU04/14); in addition to three previously sequenced whitefish; discussed below). Fish were selected from lakes Constance, Lucerne, Thun, Brienz, Biel, Neuchâtel, Zurich, and Walen which make up five separate lake-systems (Constance, Lucerne, Thun/Brienz, Biel/Neuchâtel, and Walen/Zurich; Fig. 1a; Supplementary File S1). Individuals from each whitefish species within each lake, representing the phenotypic diversity of Swiss Alpine whitefish, were sampled, including three species from Lake Constance, six from Lake Lucerne, six from Lake Thun, and four from Lake Brienz, two from Lake Biel, two from Lake Neuchâtel, three in Lake Zurich, and two in Lake Walen. In addition to these Swiss whitefish a number of outgroup individuals were also sampled, including two *Coregonus albula* (European cisco), and a number of members of the European *C. lavaretus* species complex including two Norwegian *Coregonus lavaretus*, and four samples of North German whitefish thought to be the closest relatives of the Alpine whitefish radiation members: two German *Coregonus holsatus* (from Lake Drewitz) and two German *Coregonus maraenoides* (from Lake Schaal).

The whitefish species we sampled spanned a range of six different ecomorphs that differ in their morphology, including body length, depth, and feeding morphology, as well as spawning depth and time, and diet (sampled species in each lake and the ecomorphs to which these species belong were plotted according to their distribution; Fig. 1a; Fig. 1b). Species in this study were assigned to each ecomorph based on their phenotype by whitefish taxonomic experts and co-authors Oliver M. Selz and Ole Seehausen. The ‘Balchen’ whitefish ecomorph is characterised by large bodied shallow spawning species which predominantly feed on benthic macroinvertebrates. Conversely, the ‘Albeli’ ecomorph is characterised by small species which spawn deeper (intermediate depth to very deep) and feed on zooplankton in the pelagic zone of lakes. The third ecomorph is the ‘Felchen’ type, which grow to larger sizes than the ‘Albeli’ ecomorph but not as large as the ‘Balchen’, feed on zooplankton, and feed and spawn from an intermediate depth to very deep. In addition to these three widespread ecomorphs are three less-widespread ecomorphs which occur in three or fewer lake-systems. These include two variations of profundal ecomorphs, a ‘benthic-profundal’ species, *C. profundus* from Lake Thun (an additional, now extinct, ‘benthic-profundal’ species *C. gutturosus* was also once present in Lake Constance), which have few gill-rakers but spawn at intermediate to great depth and a ‘pelagic-profundal’ species, *C. nobilis* in Lake Lucerne, which spawn deep but have a high number of gill-rakers. The final ecomorph we sampled were the ‘large-pelagic’, and included the species *C. wartmanni* from Lake Constance, *C. acrinasus* from Lake Thun, and *C. suspensus* from Lake Lucerne, which, although they are large bodied, have a high gill-raker count and feed predominantly on zooplankton. *C. wartmanni* has a well described pelagic spawning behaviour, while the other two ‘large-pelagic’ species are so far less well characterised in that respect. A full breakdown of the fish included in this study, their gill-raker counts, standard-length measurements, and the ecomorph assignment of each species can be seen in Supplementary File S1.

DNA for each individual was extracted from either fin or muscle tissue from each fish that had been stored at −80 °C using Qiagen DNeasy extraction columns, quantified using a Qubit 2.0, and run on a 1% agarose gel to assess DNA quality. DNA was then sequenced on the Illumina NovaSeq 6000 with a 550bp insert size (Next Generation Sequencing Platform, University of Bern). To this data, we added Illumina HiSeq 3000 data sequenced from one *Coregonus sp*. “Balchen” (ENA accession: GCA_902810595.1; now re-classified as *C. steinmanni* ^31^) from Lake Thun (Switzerland) that was previously used to polish and validate the Alpine whitefish reference genome assembly ^48^.

### Genotyping and loci filtering

After sequencing, all fastq files were quality checked using FastQC ^65^ before being mapped to the *Coregonus sp*. “Balchen’’ Alpine whitefish reference genome (ENA accession: GCA_902810595.1; ^48^; with additional un-scaffolded contigs (https://datadryad.org/stash/dataset/doi:10.5061/dryad.xd2547ddf) to ensure accurate mapping) using bwa-mem v.0.7.17 ^66^ changing the ‘r’ setting to 1 to allow more accurate, albeit more time-consuming, alignment. Mosdepth v.0.2.8 ^67^ was used to calculate mean sequencing coverage from the BAM files for each of the 97 individuals which ranged from 15.32x to 41.69x (an additional two individuals were added to this dataset after genotype calling discussed below). Picard-tools (Version 2.20.2; http://broadinstitute.github.io/picard/) was then used to mark duplicate reads (MarkDuplicates), fix mate information, (FixMateInformation) and replace read groups (AddOrReplaceReadGroups). Genotypes were then called across the 40 chromosome-scale scaffolds included in the *Coregonus sp*. “Balchen” Alpine whitefish assembly (ENA accession: GCA_902810595.1; ^48^) using HaplotypeCaller in GATK v.4.0.8.1 ^68^ using a minimum mapping quality filter of 30. The resulting VCF file was then filtered using vcftools v.0.1.14 ^69^ to remove indels (--remove-indels) and include biallelic loci (--min-alleles 2 --max-alleles 2) which have a minor allele count > 3 (--mac 3), no missing data (--max-missing 1), a minimum depth > 3 (--min- meanDP 3 --minDP 3), a maximum depth < 50 (--max-meanDP 50 --maxDP 50), and a minimum quality of 30 (--minQ 30), to leave 16,926,710 SNPs. Loci that fell within potentially collapsed regions of the genome assembly (as identified in De-Kayne at al. 2020) were removed using BEDTools v.2.28.0 (^70^; bedtools subtract) and any loci with duplicate IDs which were identified with PLINK v.1.90 ^71^ were removed with VCFtools ^69^ resulting in 15,841,979 SNPs. To increase our sampling of the species C*. macrophthalmus* from Lake Constance from one individual to three, we added sequencing data from an additional two individuals (Supplementary File S1). To avoid the downstream impacts of combining sequencing data from different runs (which can result from different biased nucleotide calls and introduce erroneous signals of genetic differentiation; as outlined in ^72^) we mapped these two samples as above (resulting in a mean genome-wide coverage of 9.32x and 16.58x) and called genotypes again for all samples (including the two additional C*. macrophthalmus* individuals) at each of the original 15,841,979 SNP positions. Following this genotype calling, which resulted in 15,521,925 SNPs, SNP filtering was repeated as before, leaving 14,313,952 SNPs with no missing data across the dataset of 99 individuals.

### PCA, phylogenetics, and admixture analysis

PLINK v.1.90 ^71^ was used to produce a genomic PCA of all 91 Alpine whitefish genomes with the aim of understanding how each of the individuals, species, and lakes were differentiated from one another. All eight outgroup individuals were removed from the full dataset of leaving only Alpine whitefish from the five lake-systems. Loci were then filtered based on linkage disequilibrium using PLINK v.1.90 (^71^; 50 kb windows with a step size of 10 bp and filtering for an R^2^ > 0.1). This resulted in 1,133,255 loci which were processed by PLINK to calculate eigenvector distances between individuals. PCAs were plotted using R ^73^.

We took a phylogenetic approach to understand the relationships between each of the Alpine whitefish species we sampled. First, the full VCF file was thinned to include only SNPs which were 500bp apart using VCFtools (^69^; --thin 500). The thinned SNP dataset containing 2,039,744 SNPs was then filtered using bcftools (part of SAMtools v.1.8 ^74^; bcftools view -i ‘COUNT(GT=“RR”)>0 & COUNT(GT=“AA”)>0’) to leave only SNPs that were present at least once in our dataset as homozygous for the reference allele, and homozygous for the alternative allele, as required by RAxML. This reduced the dataset to 1,692,559 SNPs. This filtered VCF file was then converted to a PHYLIP file using vcf2phylip v.2 ^75^ before RAxML v.8.2.12 ^76^ was run with the ASC_GTRGAMMA substitution model (-m ASC_GTRGAMMA --asc-corr=lewis, -k -f a) with 100 bootstraps and specifying the *C. albula* samples as outgroups to produce the maximum likelihood tree. The phylogenetic tree, excluding the long node to *C. albula*, was then plotted using Figtree v.1.4.4 ^77^.

The same linkage-pruned dataset of 1,133,255 SNPs that was used to produce the full PCA was used to calculate admixture proportions. The .bed file from PLINK resulting from the PCA was analysed using admixture v.1.3.0 ^78^ to estimate admixture for values of K between 2 and 14 specifying 20 cross validations (--cv=20). As the CV error increased with the range of K that we tested (Supplementary Fig. 2, we selected the K which helped to resolve the lake-systems and deep clade splits best, K=7, and plotted admixture barplots in R (additional admixture plots for K=2-K=10 can be found in Supplementary Fig. 3).

### Outlier scans

To identify the degree of genetic parallelism between ‘Balchen’ and ‘Albeli’ whitefish species from across the radiation, we subsetted 24 individuals representing three ‘Balchen’ species and three ‘Albeli’ species from four of the lakes we sampled: Lake Brienz, Lake Lucerne, Lake Walen and Lake Neuchâtel out of our full 99 individual dataset. ‘Albeli’ species included *C. candidus, C. albellus*, *C. muelleri*, and *C. heglingus* (for lakes Neuchâtel, Brienz, Lucerne, and Walen), and ‘Balchen’ species included *C. palea, C. alpinus, C. litoralis*, and *C. duplex* (for lakes Neuchâtel, Brienz, Lucerne, and Walen). To first confirm the independent evolution of each ‘Balchen’ and ‘Albeli’ species-pair within each of these four lakes, as indicated by the phylogeny, F4 statistics were calculated across a four-taxon tree (as used in ^42^), allowing us to estimate the degree of correlated allele frequencies between ‘Balchen’ and ‘Albeli’ individuals within and between lake-systems. First, loci were pruned based on linkage disequilibrium using the script ldPruning.sh (https://github.com/joanam/scripts/raw/master/ldPruning.sh), resulting in 1,315,105 SNPs. Then the script plink2treemix.py (from https://speciationgenomics.github.io/Treemix/) was used to convert data into the treemix format before F4 calculations were implemented using f4.py (https://raw.githubusercontent.com/mmatschiner/F4/master/f4.py). We calculated F4 for two different topologies, placing ‘Balchen’ and ‘Albeli’ species from all pairwise combinations of the four lakes on a four-taxon tree. In the first four taxon tree ((A,B),(C,D)) we placed sympatric ‘Balchen’ and ‘Albeli’ species from a first lake as A and B, and ‘Balchen’ and ‘Albeli’ species from a second lake as C and D. In this context the resulting F4 (F4^1^) represents the correlated allele frequency between A or B and C or D that would indicate introgression, or in our case, representative of a single evolution of ‘Balchen’ and ‘Albeli’ followed by sorting into lakes. We then calculated F4 where allopatric ‘Balchen’ species from two different lakes were placed as A and B and allopatric ‘Albeli’ species from the same two lakes as C and D (F4^2^). F4 in this second arrangement represents the correlated allele frequencies of sympatric species, again between A or B and C or D. Where F4^1^ < F4^2^ there is stronger support for the scenario in which ‘Balchen’ and ‘Albeli’ are truly sympatric species pairs, and therefore independently originated across lakes rather than for a single origin of the two ecomorphs.

To explore whether ‘Balchen’ and ‘Albeli’ species of whitefish show a parallel genetic basis of evolution in different lakes, regardless of lake structure, we used the cluster separation score (CSS; introduced by Jones *et al*. ^43^ and the therein reported formula corrected by Miller *et al*. ^44^), a measure of genomic differentiation between individuals assigned to two groups. Here we assigned individuals from the four ‘Balchen’ species to one group and those from the four ‘Albeli’ species to another. When calculated in windows of the genome, the CSS score quantifies the genetic distance between these ecomorph groups relative to the overall genetic variance in this particular window ^43^. We calculated CSS in 50 kb windows using a custom R script (https://github.com/marqueda/PopGenCode/blob/master/CSSm.R) where the 24 whitefish individuals were split into two groups according to ecomorph (i.e. ‘Balchen’ or ‘Albeli’). A stratified permutation test which reshuffles the assignment of individuals to each of the ecomorph groups within each lake to test the statistical significance of the CSS score for each window, whilst maintaining population structure, was then carried out 100,000 times using a custom R script (https://github.com/marqueda/PopGenCode/blob/master/CSSm_permutation.R). Windows with fewer than 24 SNPs were removed (in accordance with ^44^) and outlier windows were identified based on a false discovery rate adjusted p-value cutoff of p < 0.01, using ‘fdr.level = 0.01’ in the R package ‘qvalue’(^79^; similarly to ^44^). The median CSS score across all 34,102 windows with ≥ 24 SNPs was 0.0083 and the median CSS score across all 1659 outlier windows was 0.0973. A PCA was then produced for all 91 Alpine whitefish (excluding our outgroup samples) using PLINK v.1.90 starting with only the 690,101 SNPs that fell within these 1659 CSS outlier windows. Filtering for linkage disequilibrium was carried out as above, resulting in 56,127 SNPs that were then used to determine the genomic variation between whitefish species within these genomic regions. Correlations between PC1, which separated out species, and traits (gill-raker count and standard length) were carried out using the linear model function (lm) in R.

To confirm that this pattern is not simply driven only by the inclusion of individuals used to define the outlier CSS windows, we produced a second PCA as above but excluding the original 24 individuals. In this instance, CSS PC1 was still significantly correlated with standard length (R^2^=0.2081, p=1.183×10^-4^) and gill-raker count (R^2^=0.1135, p=5.667×10^-3^ when including the outlier *C. profundus;* R^2^=0.2201, p=1.05×10^-4^ when including the outlier *C. profundus*), albeit, and unsurprisingly, to a lesser extent.

We also identified genes that were annotated on chromosome-scale scaffolds in the whitefish reference genome ^48^ which overlapped with the 1659 outlier CSS outlier windows by a minimum of 1bp using ‘bedtools intersect’ ^70^. And then used the topGO package ^80^ in R to identify significantly enriched gene ontology terms (p-values < 0.05 according to both the ‘weight’ and ‘elim’ algorithms) associated with these outlier windows (Supplementary File S2). To demonstrate that the 1800 genes that overlapped with our 1659 CSS outlier windows were not substantially longer than non-overlapping genes, we compared their length distribution to the length distribution of all 42,695 genes (Supplementary Fig. 5).

We then calculated pairwise genome-wide relative divergence between sympatric ‘Balchen’ and ‘Albeli’ species for each lake separately. Weir and Cockerham FST was calculated between ‘Balchen’ and ‘Albeli’ species in each lake after filtering out loci which had a minor allele count < 1 between the two using vcftools v.0.1.14 (^69^; --weir-fst --mac 1) specifying a window size of 50 kb. Windows with fewer than 10 SNPs were removed. The mean F_ST_ of all windows along the genome was then calculated for each species-pair to determine the total extent of differentiation between sympatric ‘Balchen’ and ‘Albeli’ species. To identify regions of the genome which underpin the phenotypic contrast between ecomorphs we identified the top percentile of most differentiated windows in each lake and species-pair using R and those outlier windows which were shared between two or more species pairs were noted. As with CSS outlier windows, genes that overlapped with the top 1% outlier windows from each of the four species pairs were identified using ‘bedtools intersect’ ^70^. KEGG orthology was identified for 28,673 of the 46,397 annotated genes in the whitefish *Coregonus sp*. “Balchen” assembly using BlastKOALA (https://www.kegg.jp/blastkoala/; using the taxon id 861768 and selecting the genus_eukaryotes database) and as a result the genes and KEGG orthology terms that overlapped with each of the F_ST_ outlier windows, and genes overlapping with these windows, for each of the four species-pair comparisons were identified. For each species-pair the KEGG gene pathways that were associated with KEGG orthology terms associated with lake-specific F_ST_ outlier windows were also identified using the KEGG orthology database (https://www.kegg.jp/kegg/ko.html). The genes, KEGG orthology terms and KEGG gene pathways that were associated with each species-pair-specific set of F_ST_ outlier windows were then compared to identify any features that were associated with ‘Balchen’-’Albeli’ differentiation across all lake-systems. Full protein sequences for genes associated with the shared KEGG orthology terms K07526 (augustus_masked-PGA_scaffold11__203_contigs__length_63881516-processed-gene-394.0 and maker-PGA_scaffold9__196_contigs__length_60468309-snap-gene-345.2) and K12959 (maker-PGA_scaffold11__203_contigs__length_63881516-snap-gene-396.10 and maker-PGA_scaffold9__196_contigs__length_60468309-snap-gene-342.13) were BLASTed using blastp (https://blast.ncbi.nlm.nih.gov/Blast.cgi?PAGE=Proteins) and the resulting best hits, those with the highest E-value and an annotated gene name in a salmonid species were noted (Supplementary File S3).

### Genome-wide association mapping

To identify the genetic basis of gill-raker variation across the Alpine whitefish radiation we used a mixed model approach implemented in EMMAX ^81^(as in ^14^). First, EMMAX was used to produce a Balding-Nichols kinship matrix between all 90 Alpine whitefish samples for which we had gill-raker counts using ‘emmax-kin’ using only the 9,120,498 SNPs that were polymorphic within the Alpine whitefish radiation. We then used EMMAX to calculate the association of each SNP marker with gill-raker count for each SNP. Two significance thresholds were determined. A strict Bonferroni multiple testing p-value threshold was calculated using the total number of SNPs tested: −log10(0.05/9120498) = 8.26, in addition to an LD-considerate threshold of −log10(0.05/4536915) = 7.96, which was calculated by removing linked markers (R^2^ > 0.95) in 50 kb sliding windows across the genome using PLINK ^71^. One SNP on WFS23 had an association above the LD-considerate threshold and the allele frequencies within each of the six ecomorph groups was calculated for this SNP using vcftools --freq on each subset of ecomorphs separately (Fig. 2e; in addition to each ecomorph within each lake separately; Supplementary Fig. 8). The gene that overlapped with this SNP was identified with BEDTools ^70^ and the full protein sequence from the gene that overlapped with the SNP (maker-PGA_scaffold22__199_contigs__length_52020451-snap-gene-302.9) was BLASTed using Ensembl TBLASTN against the Atlantic Salmon, Rainbow Trout, Brown Trout and Coho Salmon genomes, hitting with high confidence against the ectodysplasin-A receptor (*edar*) gene (E-value 1e-20; ID% 97.62 in Brown Trout fSalTru1.1; ENSSTUG00000036900 and E-value 7e-20; ID% 100 in Atlantic Salmon ICSASG_v2; ENSSSAG00000053655). The variance in gill-raker count across our samples explained by the most significantly associated SNP was calculated using the equation: PVE = ((2*(beta^2)*MAF*(1-MAF))/(2*(beta^2)*MAF*(1-MAF)+(se_beta^2)*2*N*MAF*(1-MAF))) where N = the sample size (90), se_beta = the standard error of effect size of the SNP, beta = SNP effect size, and MAF = SNP minor allele frequency (from the Supplementary Information S1 associated with ^82^).

This EMMAX association mapping was repeated using sex as a binary trait for 90 Alpine whitefish individuals. The most substantial associated peak was observed on WFS04. As above, genes that overlapped with these SNPs were identified with BEDTools ^70^ and the protein sequence from the single gene that overlapped with this peak of SNPs on WFS04, maker-PGA_scaffold3__454_contigs__length_92224161-snap-gene-551.2, was BLASTed using Ensembl TBLASTN against the Atlantic Salmon, Rainbow Trout, Brown Trout and Coho Salmon genomes, however, no annotated genes were hit with high confidence using this approach.

### F-branch statistics

To calculate excess allele sharing across the dataset, and test whether species of the less-widespread ecomorphs with unique trait combinations (i.e. combinations of traits that contrast with the direction of correlation among combinations of traits seen in the widespread ecomorphs) have evolved as a result of gene flow between lake-systems, we used the f-branch statistic *f_b_*(*C*) as calculated by the package Dsuite ^83^ as in ^84^. First, a simplified version of the full RAxML phylogenetic tree was prepared. To make use of the multiple samples per species in our dataset and get robust estimates of excess allele sharing both within and between lake-systems, collapsed nodes in the phylogenetic tree using the R package ‘ape’ ^85^ where possible. Individuals which looked like potential F1 hybrids as indicated by close to 50/50 splitting in the admixture analysis or were placed discordantly in our genome-wide PCA and phylogeny (including the *C. alpinus* 0129 and *C. zuerichensis* 099) and individuals which did not sit in the same clade as other individuals of the same species in the same lake-system were kept separated so as to not skew species-wide estimates of excess allele sharing from single, potentially recent introgression events, and thus not included in node collapsing. Nodes were then collapsed, and the individuals within that clade assigned as a single tree tip, if all individuals within the clade belonged to the same species or species of the same ecomorph from a single lake or, where possible, single lake-system (excluding potential F1 individuals). All outgroup individuals in the tree were collapsed into a single outgroup tip. Dsuite ^83^ was then run specifying Dtrios, DtriosCombine, and finally Fbranch, each time specifying the collapsed tree. Dsuite was used to first calculate f4 admixture ratios f(A,B;C,O) across the dataset where combinations of taxa fit the necessary relationship ((A, B), C) in our phylogenetic tree, with the 8 non-Alpine whitefish set as the outgroup. The f-branch statistic *f_b_*(*C*) was then calculated from these f4 statistics using the phylogenetic tree to identify excess allele sharing between any taxa into any other taxon or node in the phylogeny. *f_b_*(*C*) is particularly powerful for complex systems such as the Alpine whitefish radiation since, unlike Patterson’s D, it provides branch-specific estimates of excess allele sharing, meaning that specific instances of gene flow do not skew excess allele sharing estimates across multiple nodes or branches, providing a phylogenetically-guided and robust estimate of excess allele sharing ^84^. Significant instances of excess allele sharing were identified by calculating a stringent Bonferroni multiple-testing significance threshold, which involved dividing the p-value threshold of p < 0.01 by the number of cells in the f-branch matrix for which *f_b_*(*C*) could be calculated (1910) and converting this to a Z-score using R. All cells with Z-scores higher than this threshold i.e. Z > 4.41 represented significant excess allele sharing between taxa in the tree and were indicated as such.

## Data availability

The raw sequencing files will become accessible on SRA upon publication (and the appropriate SRA sample codes added to Supplementary File S1) and additional source data (genotype file and corresponding metadata file along with figure-specific data) will be deposited on the Eawag research data institutional collections (https://doi.org/10.25678/0005S0) upon publication.

## Code Availability

Scripts for all analyses are available on GitHub: https://github.com/RishiDeKayne/Alpine_whitefish_WGS.

## Acknowledgments

We thank Anna Feller and Adam Ciezarek for comments on earlier versions of the manuscript. We acknowledge Verena Kälin for the whitefish illustrations. The data produced and analysed in this paper were generated in collaboration with the Next Generation Sequencing Platform, University of Bern, and the Genetic Diversity Centre (GDC), ETH Zurich. This work was supported by the Swiss National Science Foundation (SNSF project 31003A_163446/1 awarded to P.G.D.F.)

## Author contributions

R.D.K, O.S., P.G.D.F. conceived and designed the project. O.M.S. and O.S. selected whitefish individuals for sequencing. O.M.S. measured whitefish morphological traits. R.D.K, supported by P.G.D.F., carried out molecular lab work, data processing and analysis, with help from D.A.M. for CSS calculations. R.D.K. and P.G.D.F., with support from O.M.S., D.F., O.S. interpreted the results and wrote the manuscript. All authors contributed to and revised the final version of the manuscript.

## Materials & Correspondence

Correspondence and material requests should be addressed to P.G.D.F.

