## Supplementary Information for "Hybridization and a mixture of small and large-effect loci facilitate adaptive radiation"

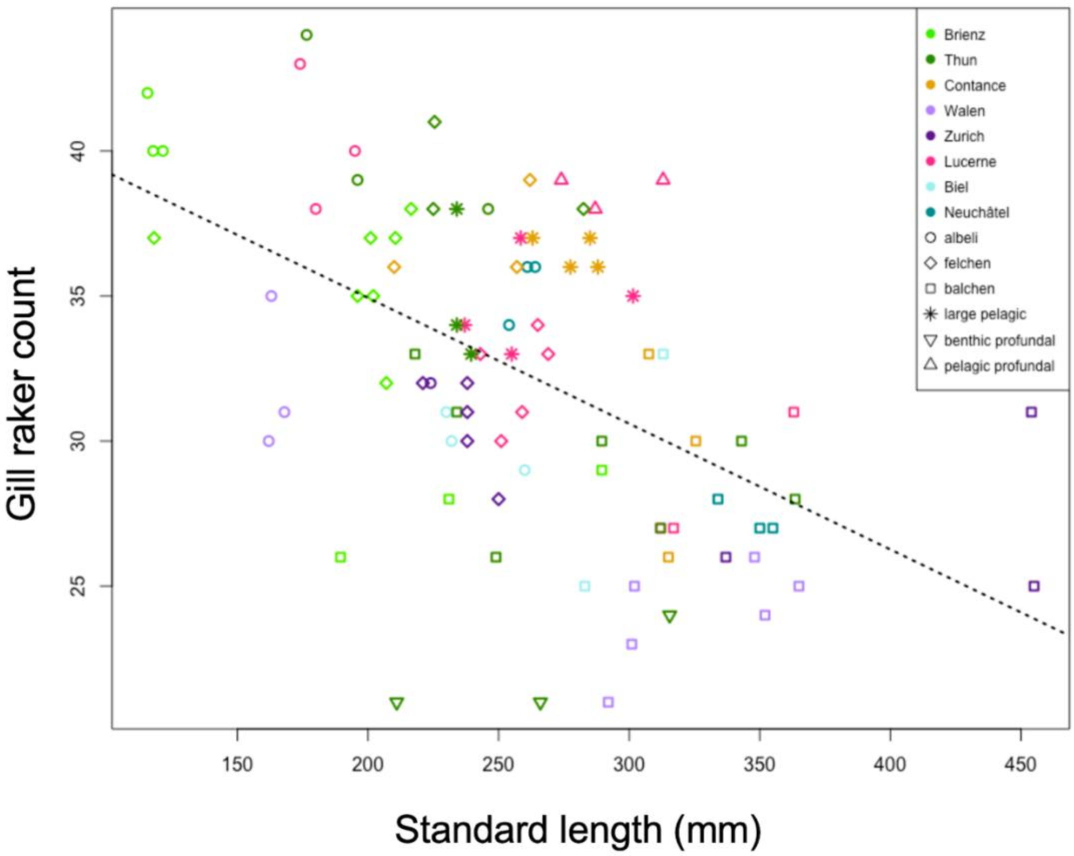

Supplementary Figure 1. Standard length and gill raker count of whitefish were correlated across the dataset (black line indicates linear regression; $R^2$=0.2767, p=1.2x$10^{-7}$). 'Albeli' species (circles) tend towards high gill raker counts and smaller standard length, and 'Balchen' species (squares) tend towards lower gill raker counts and larger standard lengths.

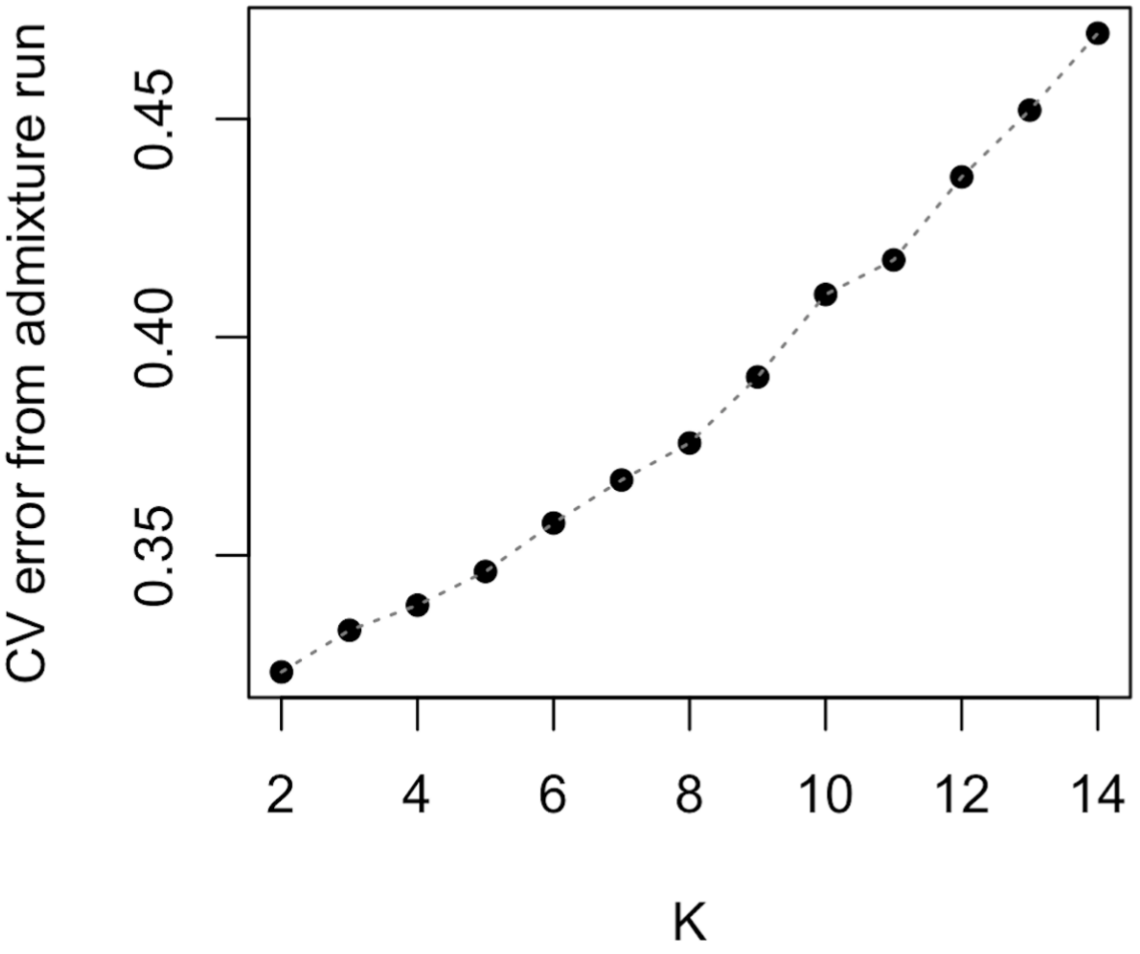

Supplementary Figure 2. CV error increased with the number of populations (K) for each admixture run. K=7 was selected for plotting since it helped identify lake-specific differences.

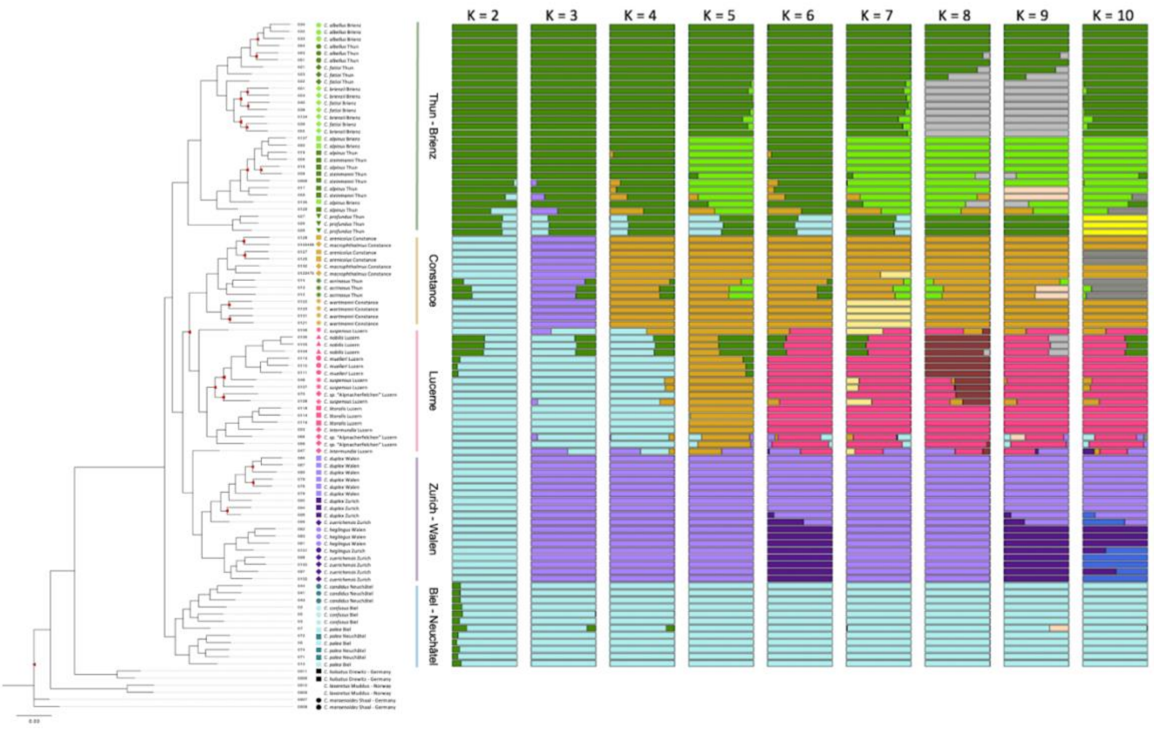

Supplementary Figure 3. Our maximum likelihood RAxML phylogeny with admixture analysis where the number of populations (K) ranges from 2 to 10.

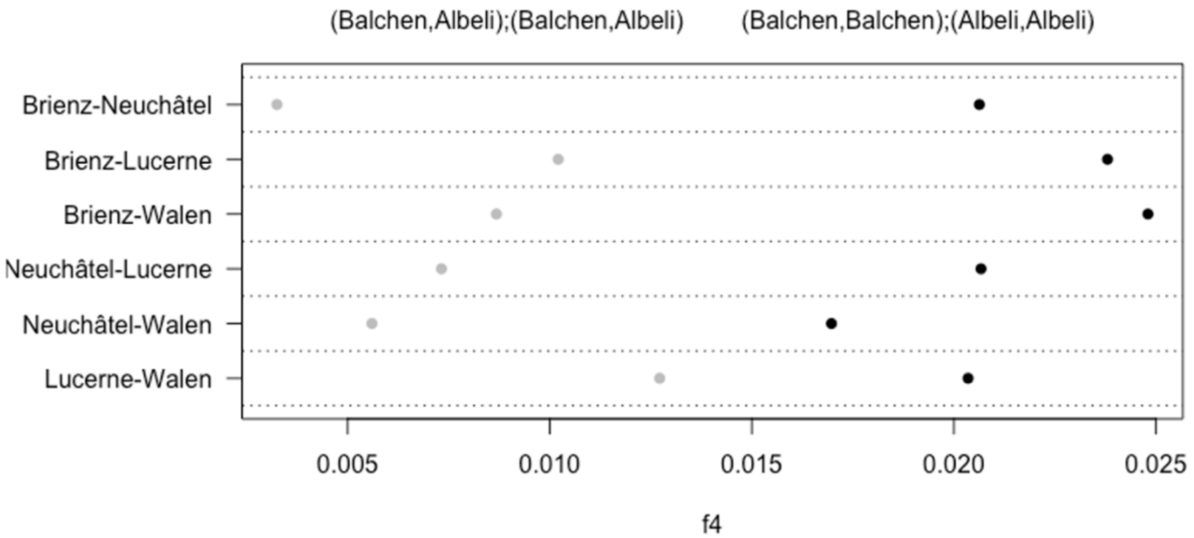

Supplementary Figure 4. F4 statistics calculated for topologies where ‘Balchen’ and ‘Albeli’ for each of the lakes Brienz, Neuchâtel, Walen, and Lucerne where either sister to one another i.e. sorted by lake (grey), or where species of the same ecomorph are sympatric to one another regardless of lake (black). f4 statistics of the topology inconsistent (right, black) with the phylogenetic tree of the adaptive radiation (Fig. 1) are higher than f4 statistics consistent with its topology (left, grey), supporting a scenario in which ‘Balchen’ and ‘Albeli’ ecomorphs evolved independently in parallel in each pre-Alpine lake (see Methods for more details).

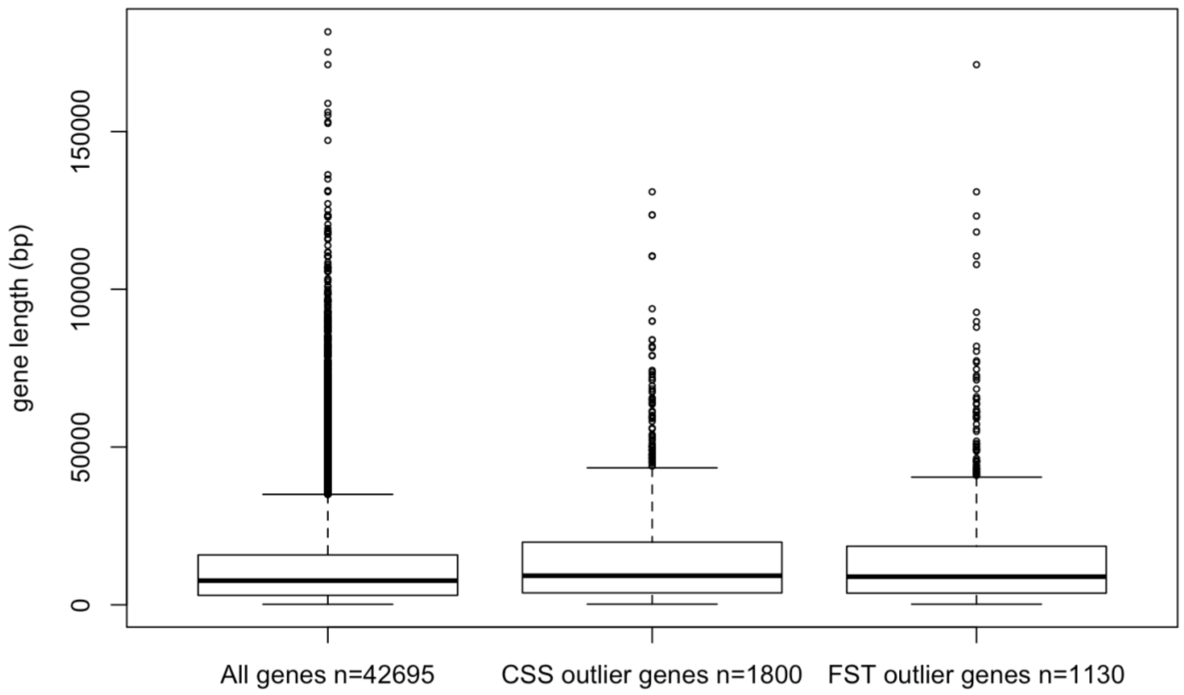

Supplementary Figure 5. Boxplot showing that the 1800 genes that overlapped with our CSS outlier windows and 1130 genes that overlapped with $F_{ST}$ outlier windows did not demonstrate a substantial skew in length compared to the full set of 42,695 annotated genes. The midline of the box represents the median gene length for each gene set, the box represents the third and first quantile of the distribution. The uppermost whisker extends to 1.5x the interquartile range, with lengths beyond this represented by outlier points, and the lowermost whisker represents the shortest gene lengths in each set.

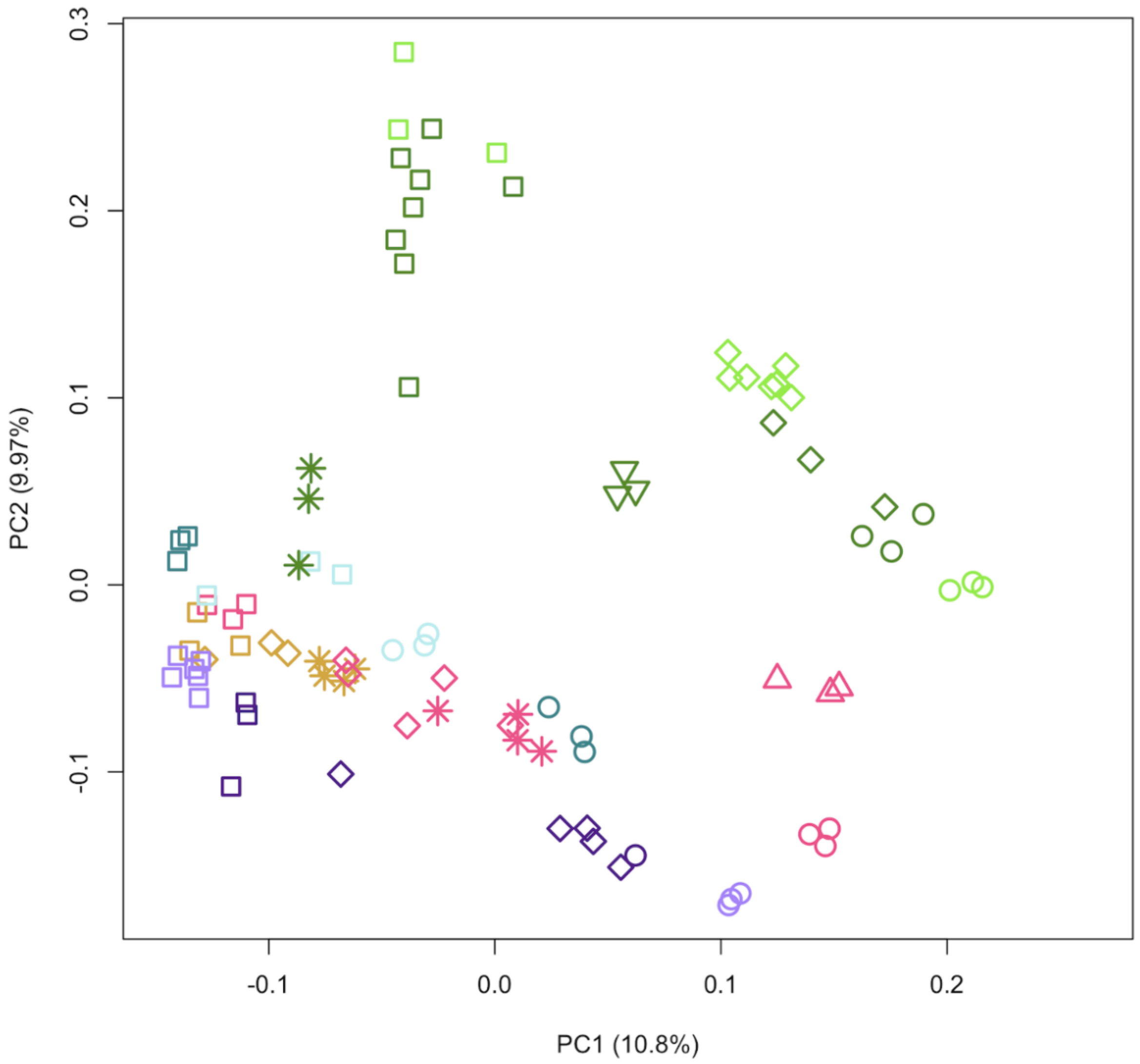

Supplementary Figure 6. In our PCA including only linkage-filtered SNPs within the 1659 CSS outlier windows PC1 separates out species and PC2 separates out the Thun/Brienz lake-system from all four other lake-systems. Symbols are used as in Fig. 1.

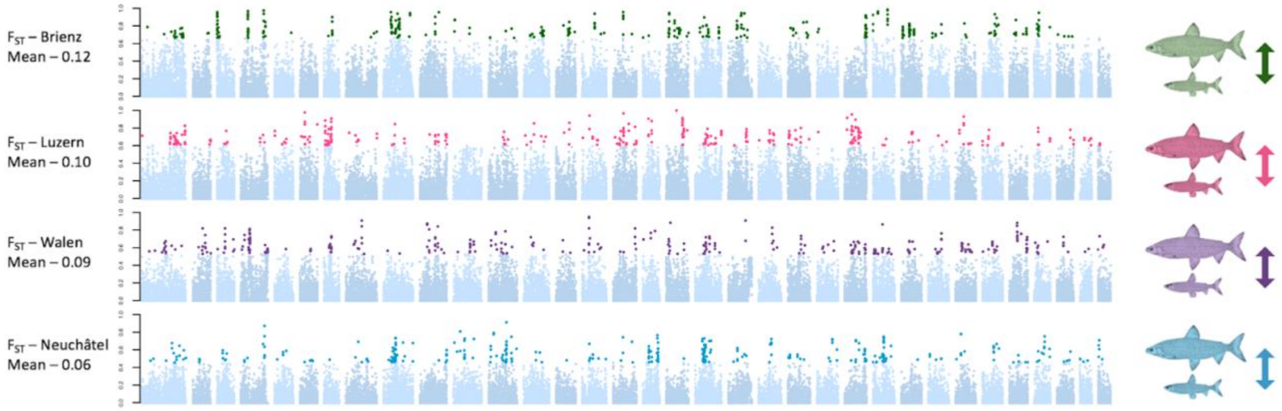

Supplementary Figure 7. Pairwise $F_{ST}$ scans (50 kb windows) for 'Balchen' and 'Albeli' species from Lakes Brienz (a), Lucerne (b), Walen (c), and Neuchâtel (d). Mean genome-wide $F_{ST}$ is shown for each pairwise comparison.

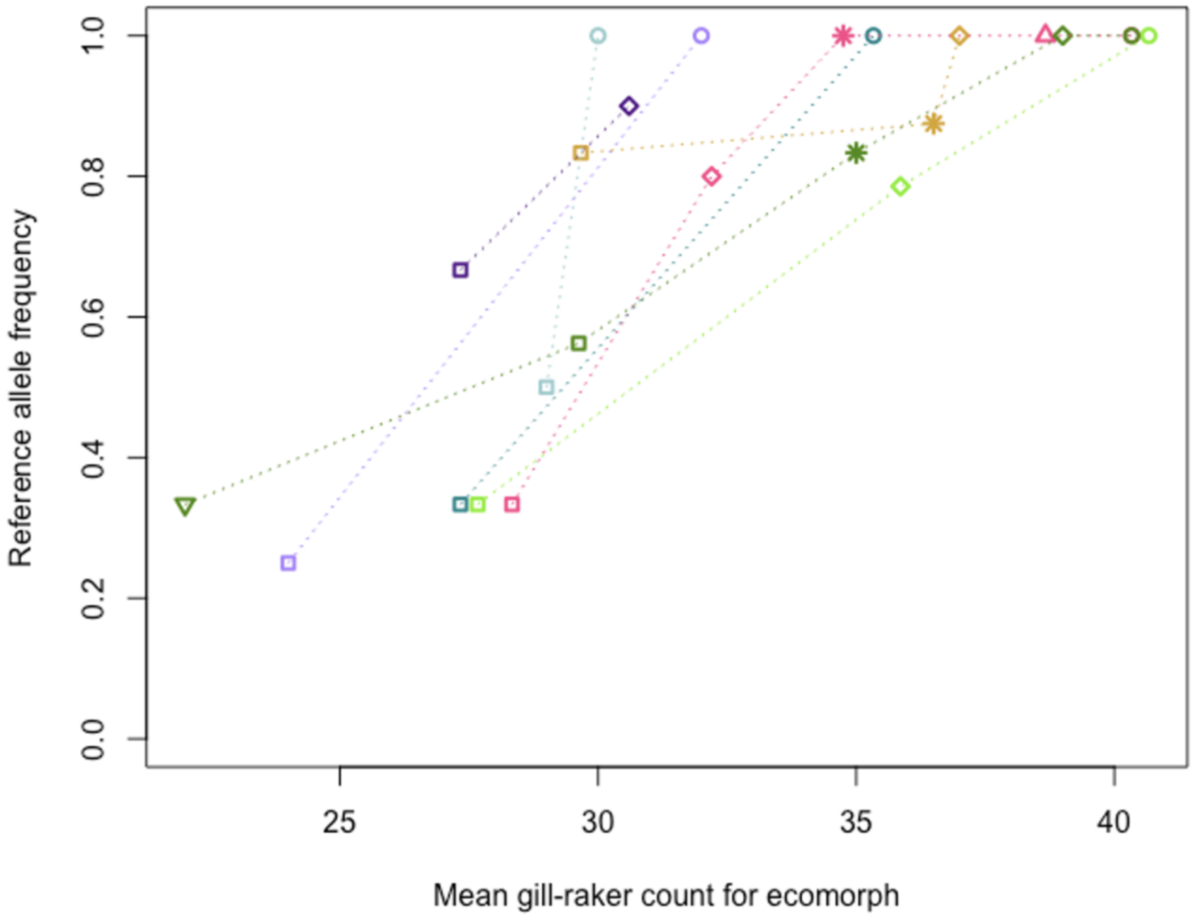

Supplementary Figure 8. Gill-raker count across species within each lake correlates to reference allele frequency for the significantly associated peak on WFS23. The reference allele frequency for each species split by lake and plotted against mean gill-raker count for those species individuals by lake. Only species with >1 individual were plotted (resulting in the exclusion of the one *C. heglingus* individual from lake Zurich). Symbols indicate ecomorph assignment and are used as in Fig. 1.

Supplementary Table S1. Full results of statistical analysis of regressions between PC1 (calculated from loci within CSS windows) and gill raker count and standard length. Regressions of phenotypes across dataset.

| | | | | |
|---|---|---|---|---|
| CSS PC1 vs. Gill-raker count | | | | |
| Lake system | DF | $R^2$ | P | significance $p < 0.001$ = *** / $p < 0.01$ = ** / $p < 0.05$ = * / $p > 0.05$ = NS |
| All | 88 | 0.3921 | 4.10E-11 | *** |
| All no outlier *C. profundal* | 85 | 0.5107 | 7.63E-15 | *** |
| Luzern | 16 | 0.8113 | 3.47E-07 | *** |
| Brienz/Thun including outlier *C. profundus* | 31 | 0.3871 | 1.11E-04 | *** |
| Brienz/Thun no outlier *C. profundus* | 28 | 0.6051 | 4.22E-07 | *** |
| Zurich/Walen | 16 | 0.7167 | 9.43E-06 | *** |
| Biel/Neuchâtel | 9 | 0.6857 | 0.001645 | ** |
| Constance | 8 | 0.3703 | 0.06194 | NS |
| CSS PC1 vs. Standard length | | | | |
| Lake system | DF | $R^2$ | P | |
| All | 88 | 0.498 | 8.06E-15 | *** |
| Luzern | 16 | 0.322 | 0.01405 | * |
| Brienz/Thun | 30 | 0.3547 | 0.0003233 | *** |
| Zurich/Walen | 16 | 0.6925 | 1.84E-05 | *** |
| Biel/Neuchâtel | 10 | 0.6738 | 0.001067 | ** |
| Constance | 8 | 0.1215 | 0.3236 | NS |
| CSS PC1 vs. Gill-raker count (excluding original 24 samples) | | | | |
| All | 64 | 0.1135 | 0.005667 | ** |
| All no outlier *C. profundus* | 61 | 0.2201 | 0.000105 | *** |
| CSS PC1 vs. Standard Length (excluding original 24 samples) | | | | |
| All | 64 | 0.2081 | 0.0001183 | *** |
| GRC vs. Standard length (2 individuals with missing values excluded) | 87 | 0.2767 | 1.20E-07 | *** |

Supplementary Table S2. Details of genes, unique KEGG orthology terms, and KEGG pathways associated with outlier windows from across all four independent lake FST comparisons between 'Balchen' and 'Albeli' species (i.e. found at least once) and the numbers of each found in all four comparisons.

| | Total present at least once across all four lakes | Number shared across all four lakes |
|---|---|---|
| Genes overlapping with top 1% $F_{ST}$ outlier windows | 1130 | 0 |
| Unique KEGG orthology terms from these genes | 660 | 2 |
| KEGG pathways from these KEGG orthology terms | 315 | 111 |
